# Widespread decoupling between abundance and genetic diversity, and strong local genetic structuring in marine unicellular eukaryotes

**DOI:** 10.64898/2026.07.16.738895

**Authors:** María Martínez-Ríos, Pablo G. Peña, Iñaki Ruiz-Trillo, Enrique Lara, Rubén González‐Miguéns

## Abstract

How abundance, connectivity, and genetic diversity covary across space remains a central question in evolutionary biology. Using a global environmental DNA metabarcoding dataset targeting mitochondrial cytochrome c oxidase subunit I (COI), we examined relationships among metabarcoding-derived abundance, nucleotide diversity, and spatial genetic connectivity across marine eukaryotes. We found a widespread but incomplete decoupling between abundance and genetic diversity, particularly in unicellular lineages. By contrast, distance-decay patterns in abundance, diversity, and connectivity were broadly similar across eukaryotes and more strongly associated with cellularity. Focusing on the Iberian Peninsula, we reconstructed locality-centred haplotype networks to quantify the spatial organization of intraspecific genetic variation. Unicellular lineages showed stronger local genetic structuring, whereas multicellular organisms exhibited clearer large-scale biogeographic partitioning. Overall, our results reveal shared biogeographic organization across eukaryotes while demonstrating that abundance-based biodiversity metrics alone cannot fully capture evolutionary and demographic processes, especially in microbial organisms.

## 1. Introduction

Demographic processes and their spatial structuring are central to evolutionary theory, as they determine how mutation, drift, gene flow, and selection shape genetic variation (Bradburd & Ralph 2019; Lewontin 1974). Under neutral expectations, genetic diversity reflects the combined effects of mutation rate and long-term effective population size (Ne), and is therefore expected to increase with population size and connectivity (Ellegren & Galtier 2016). However, comparative studies across multicellular eukaryotes have revealed weak and taxon-dependent relationships between genetic diversity and proxies of census population size, a pattern underlying Lewontin’s paradox of variation (Buffalo 2021; Charlesworth & Jensen 2022; Leffler *et al*. 2012; Romiguier *et al*. 2014). Whether this apparent decoupling between abundance and genetic diversity extends to microbial eukaryotes remains largely unresolved.

Protists, which are predominantly unicellular, represent the majority of eukaryotic diversity (Burki *et al*. 2021; de Vargas *et al*. 2015), and provide a critical test of whether abundance, connectivity, and genetic diversity are coupled in the same way across eukaryotes. Despite often reaching extraordinarily large census population sizes, some planktonic protists show unexpectedly modest levels of genetic diversity (Filatov 2019; Krasovec *et al*. 2020; Krueger-Hadfield *et al*. 2014). This apparent mismatch suggests a potential breakdown of the classical relationship between abundance and population size. Several non-mutually exclusive mechanisms have been proposed to explain this discrepancy, including complex life cycles with alternating reproductive modes (Sundqvist *et al*. 2018), dormancy (Rengefors *et al*. 2017), and episodic blooms (Krueger-Hadfield *et al*. 2014) acting on large populations. Whatever the reasons might be, it remains to be tested how far these observations can be extended to all unicellular eukaryotes, and, in line, how critically the situation changes with respect to multicellularity. A potential gap between protists and multicellular organisms would have important implications because, if abundance and genetic diversity are partially decoupled, classical biodiversity metrics may fail to capture the evolutionary and demographic processes shaping the distribution of unicellular eukaryotes (McGaughran 2015; Overcast *et al*. 2021). Notably, a general framework linking abundance, connectivity, and genetic diversity across eukaryotes should be revised.

In this study, we address this gap by explicitly testing the relationships among genetic diversity, relative abundance, and spatial genetic connectivity across marine eukaryotes. Using mitochondrial cytochrome oxidase subunit I (COI) metabarcoding data, we examine the relationship between nucleotide diversity, spatial genetic connectedness, and a sequence-derived proxy of relative abundance across marine eukaryotes. To achieve this, we developed a network-based framework that translates haplotype relationships into locality-level measures of genetic connectivity, enabling the integration of spatial structure and intraspecific variation within a unified analytical approach (Figure 1). We first assessed global patterns across eukaryotic phyla using the global eKOI metabarcoding dataset (González-Miguéns *et al*. 2025). We then focused on marine systems surrounding the Iberian Peninsula, a well-characterized biogeographic transition zone between the North Atlantic Ocean and the Mediterranean Sea (Patarnello *et al*. 2007; Spalding *et al*. 2007). By combining correlation analyses, spatial decay models, and haplotype-network structure, we test whether classical expectations linking abundance and genetic diversity hold across major eukaryotic lineages, or whether these relationships become decoupled in microbial eukaryotes. Furthermore, we also compared how distance affects abundance, connectivity, and intraspecific diversity in unicellular and multicellular eukaryotes, as well as across taxonomic groupings. More broadly, our study also provides a scalable framework to infer population-level processes from metabarcoding data, offering new insights into whether fundamental expectations of population genetics generalize across marine eukaryotes.

**Figure 1.**
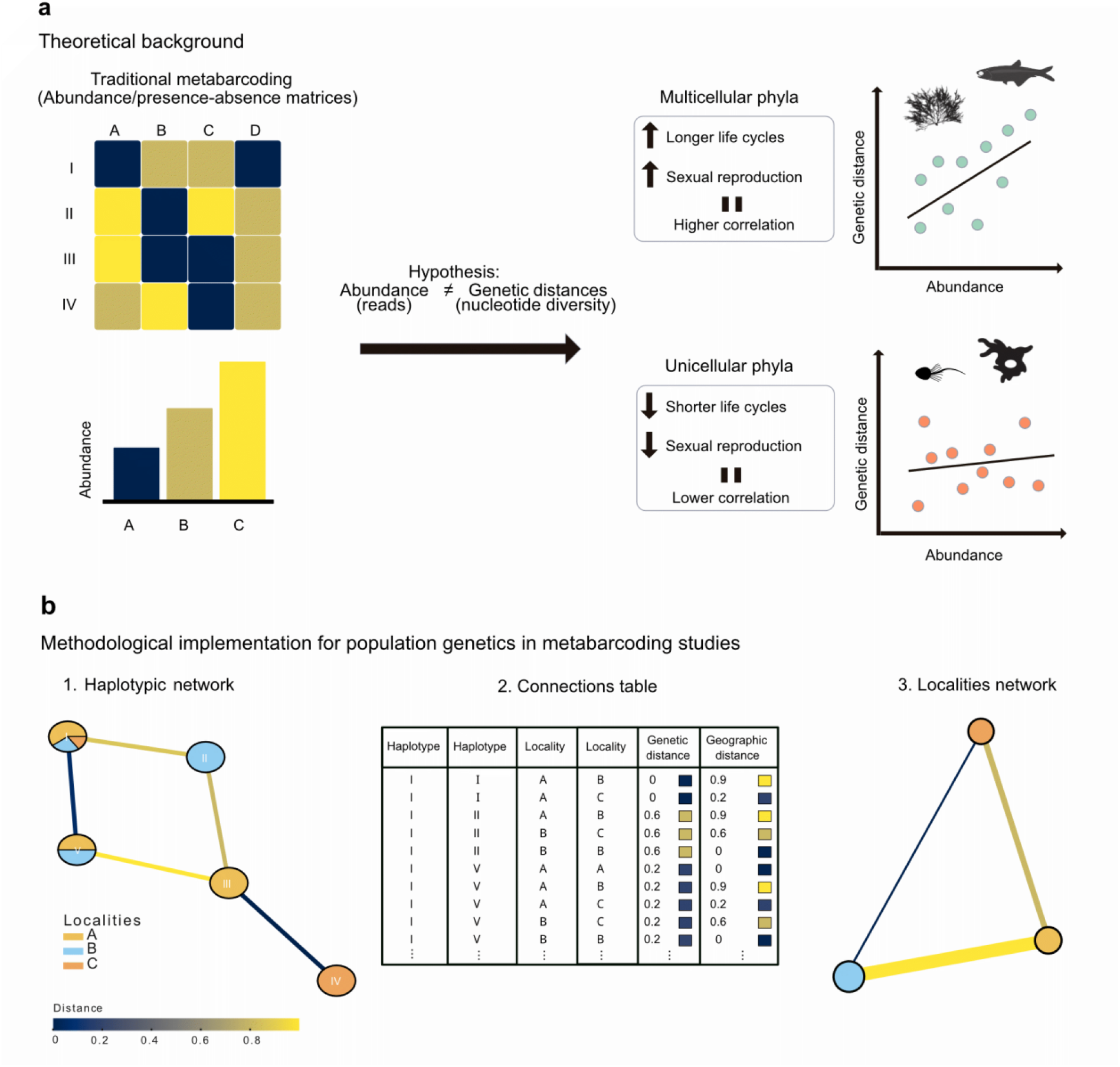
Schematic representation of the study’s hypothesis and methodological framework. **a**, Theoretical background: conceptual diagram illustrating the hypothesis that abundance and nucleotide (genetic) diversity are more strongly correlated in multicellular organisms, which typically have longer life cycles and sexual reproduction, than in unicellular taxa, characterized by shorter life cycles and rapid asexual reproduction. **b**, Methodological implementation: workflow showing the integration of haplotype-network information into metabarcoding analyses. For each OTU, haplotype networks were used to extract pairwise connections among haplotypes, generating a connection table containing genetic, geographic, or ecological distances between pairs. This information was then used to construct locality-based networks, allowing inference of spatial patterns of genetic connectivity.

## 2. Material and methods

### 2.1. Dataset formation: eKOI database ver. 2.0

To test both our original hypotheses and the proposed analytical framework, the first step was to generate a comprehensive global database suitable for these analyses. For this purpose, we used the global eKOI database (González-Miguéns *et al*. 2026), which compiles marine metabarcoding studies targeting the mitochondrial cytochrome c oxidase subunit I (COI) gene, a widely used marker for eukaryotic lineages (Hebert *et al*. 2003). We expanded this database to version 2.0 by incorporating additional metabarcoding studies, resulting in a total of 101 studies (Data S1).

Furthermore, to enhance the representativeness of the Iberian Peninsula dataset, additional samples were collected along the Atlantic and Mediterranean coasts of the Iberian Peninsula (Data S1). We collected a total of 108 samples, covering 30 coastal localities from the Mediterranean and Atlantic regions. The samples were taken from the supralittoral zone of the beaches and were fixed using LifeGuard Soil Preservation Solution (Qiagen). Total eDNA was extracted using the DNeasy PowerSoil Pro Kit (Qiagen) following the instructions provided by the manufacturer and preserved frozen (−20°C) until further processing. Then, we amplified a portion of the gene COI using the primer pair jgHCO2198 (Geller *et al*. 2013) and mlCOIintF-XT (Wangensteen *et al*. 2018). The PCR cycling profile was as follows: initial denaturation at 94°C for 3 min, followed by 35 cycles of denaturation at 94°C for 1 min, annealing at 45°C for 1 min, and extension at 72°C for 1 min, with a final extension step at 72°C for 10min. Each PCR was replicated three times, and then all replicates were pooled together to mitigate the effects of PCR biases. Amplified products were sequenced using Illumina NextSeq 2000 using a P1 600-cycle flow cell at the Fundación Parque Científico of Madrid (Spain).

The amplicon sequence variants (ASVs) of the total eKOI metabarcoding database 2.0 were then taxonomically assigned at the phylum level using the eKOI taxonomic database integrated in PR^2^ (González-Miguéns *et al*. 2025) and VSEARCH (Rognes *et al*. 2016), applying a minimum identity threshold of 84%. We grouped the ASVs per phylum and, then, we performed a multiple sequence alignment using MAFFT v7 (Katoh & Standley 2013) with default parameters. The ASVs were clustered together into Operational Taxonomic Units (OTUs) to generate molecular units. For that purpose, we used a 97% sequence similarity threshold, corresponding to a commonly applied 3% divergence cutoff in COI metabarcoding studies (González-Miguéns *et al*. 2025), using VSEARCH with the --cluster_fast algorithm.

To ensure robust population-genetic analyses and minimize potential artifacts, we applied stringent filtering criteria to retain only informative OTUs (González-Miguéns *et al*. 2026), defined as those that (i) contain at least four sequences from a minimum of two distinct locations, and (ii) include at least one pair of sequences differing by ≥ 1% (uncorrected p-distance). Then, we retained only those phyla containing at least 10 informative OTUs.

#### 2.2.1 Nucleotide diversity, relative abundance, and haplotype connections

For each informative OTU, we calculated three metrics: (i) nucleotide diversity, using the *pegas* package (Paradis 2010) in R; (ii) relative abundance, computed as the sum of ASV read counts and, subsequently, log-transformed using log_1p_(abundance) to normalize the distribution (a proxy used for abundance in metabarcoding studies (Lamb *et al*. 2019)), and; (iii) haplotype connectivity, inferred from haplotype networks constructed with the *pegas* package, where the total number of connections per OTU was log-transformed using log_1p_ (connections). Then, for the OTUs results for each phylum, we computed Spearman’s rank correlation coefficients (ρ) between nucleotide diversity, relative abundance, and genetic connectivity to evaluate the strength and significance of pairwise metric associations.

To further investigate whether the relationships among these metrics differed across major eukaryotic lineages, we conducted group-level comparisons of ρ values under two classification schemes: (i) broad taxonomic groups, including animals (Metazoa), plants (Archaeplastida), and protists/fungi; and (ii) cellular organization, separating unicellular from multicellular taxa. For each pairwise combination of variables (genetic diversity vs. relative abundance, genetic diversity vs. haplotype connectivity, and haplotype connectivity vs. relative abundance). Spearman’s ρ values was recalculated across 1,000 bootstrap resamples within each group and classification scheme. Differences among bootstrap distributions were summarized using one-way ANOVA followed by Tukey’s honestly significant difference (HSD) post hoc tests, and compact letter displays were used to indicate statistically homogeneous groups. Non-parametric Kruskal-Wallis tests followed by pairwise Wilcoxon rank-sum tests with Benjamini–Hochberg correction were additionally performed as sensitivity analyses. All analyses were conducted in R using the *rstatix* (ver. 0.7.2), *multcompView* (0.1-11), *ggpubr* (0.6.3), and *ggplot2* (Wickham 2009) packages.

#### 2.2.2 Spatial distance-decay curves

To place the previously described metrics (nucleotide diversity, relative abundance, and haplotype connectivity) in a geographic context, we examined how each metric varied with spatial separation by computing distance-decay curves. To reduce spatial redundancy and facilitate interpretation, for each informative OTU, samples located within a 5 km radius were grouped into geographic clusters based on pairwise Haversine distances using the *geosphere* package (ver. 1.5) in R. For each geographic cluster, we then calculated the same metrics as described above.

For each OTU and for each metric independently, the cluster with the highest value for a given metric was designated as the cluster centre (geographic distance = 0). Geodesic distances from each cluster to this centre cluster were then computed. Using these per-cluster values, we fitted five independent distance-decay models: (i) a linear model (LM) representing direct proportionality; (ii) an exponential decay model, fitted using nonlinear least squares (NLS); (iii) a power-law model, fitted in log-log space to capture scale-dependent relationships; (iv) a Gompertz model, representing an asymptotic sigmoidal growth function; and (v) a generalized additive model (GAM) fitted with non-parametric smooth functions to detect non-linear trends without imposing a specific functional form.

Each model was fitted using all pairwise combinations of per-cluster values of nucleotide diversity, log-transformed abundance, and log-transformed numbers of haplotype connections. Analyses were performed independently for each OTU. Model parameters were estimated using iterative optimization routines, and convergence was verified for all non-linear models. Model performance was evaluated using Akaike’s Information Criterion (AIC) and adjusted R^2^, balancing explanatory power and model complexity. Only models with stable parameter estimates were retained. All model coefficients, correlation values, and AIC scores were compiled across OTUs and summarized at the phylum level.

Then, we tested whether pairwise relationships among metrics differed significantly within each phylum, based on the OTU-level model outputs. Similarly, for each metric, we assessed differences among phyla. Permutation-based tests were applied to compare model slopes across metrics and phyla. Multiple testing correction was applied using the Benjamini-Hochberg false discovery rate (FDR) procedure. Comparative heatmaps were generated to visualize cross-phylum similarities in model performance, focusing particularly on the best-fitting model for each lineage. Asterisks denote FDR-adjusted significance levels from permutation-based pairwise tests: adjusted p < 0.05, adjusted p < 0.01 and adjusted p < 0.001. Rows and columns were ordered by hierarchical clustering to visualize cross-phylum similarities in distance-decay behaviour.

### 2.3. Geographic haplotype networks in the Iberian Peninsula

We optimize a network-locality framework that translates haplotype networks into locality-level connectivity models. We applied this approach to the Iberian Peninsula samples in the updated eKOI database. For this purpose, we first generated a subset of the dataset by retaining only the informative OTUs for each phylum present in the Iberian Peninsula and for which at least three haplotypes were detected in this region. For each OTU, haplotype networks were reconstructed and subsequently transformed into locality-based connectivity graphs, where nodes represented sampling localities and edges denoted genetic connections inferred from shared haplotypes. Specifically, for each informative OTU-specific haplotype network, we labelled connections as (i) “same” when connected haplotypes were identical, and (ii) “different” when connected haplotypes were different across localities. These connections were then compiled into a matrix that captured, for every pairwise link, both the genetic distance and geographic distance between localities in which all the analyses described above were performed.

To detect modular structure among localities, we estimated network modularity using the Louvain algorithm implemented in the *igraph* R package (Csardi & Nepusz 2005), allowing the identification of geographically coherent clusters of interconnected sampling sites. Modularity analyses were performed using the total number of connections among localities, as well as alternative weighting schemes motivated by the expectation of isolation by distance (IBD), widely debated and documented in protists. Thus, four weighting schemes were applied to represent the strength of locality-to-locality links: (i) connection count (number of shared haplotypes), (ii) inverse genetic distance (1/d), (iii) linear scaling (1 - d / max d), and (iv) a radial basis function (RBF; exp(-d/σ)), with σ optimized to maximize modularity. Independent locality clusters were obtained for each weighting approach.

To test for spatial autocorrelation in the resulting genetic structure, Moran’s I statistics were computed for each inferred locality cluster using the *spdep* R package (ver. 1.3), evaluating whether genetically similar localities were spatially aggregated or randomly distributed. P-values were adjusted for multiple testing using the Benjamini-Hochberg false discovery rate (FDR).

In addition, clusters were characterized according to Mediterranean-Atlantic composition (Patarnello *et al*. 2007) to quantify regional connectivity and isolation, providing a quantitative measure of regional connectivity and isolation. For each cluster including at least 3 localities, we calculated the percentage of localities assigned to each category, based on the oceanographic metadata assigned to each locality.

Finally, we quantified geographic distances of genetic connections using Haversine distances between linked localities implemented in the *geosphere* R package (ver. 1.5). To minimize the influence of the long-distance outliers, median distances were calculated for “all”, “same”, and “different” connections, both globally and per OTU. Statistical comparisons between the “same” and “different” connection sets were performed using Welch’s t-tests (unequal variances) and Hedges-corrected Cohen’s d as a measure of effect size using the *rstatix* R package (ver. 0.7.2).

## 3. Results

### 3.1. Marine metabarcoding dataset

The final eKOI ver. 2.0 database comprises 1,095,897 amplicon sequence variants (ASVs) and 362,781,685 reads, distributed across 1,097 unique localities and 4,659 samples (see Data S1 and Data S2 in Supporting Information). For the remaining analyses, we selected only informative OTUs, defined as those present in at least two localities, comprising at least four ASVs, and showing ≥1% molecular divergence (see Materials and Methods). To avoid concluding poorly represented taxonomic groups, we further restricted analyses to phyla containing at least 10 informative OTUs. The resulting dataset includes 1,567 informative OTUs spanning 20 eukaryotic phyla (Data S3).

### 3.2. Testing the abundance-diversity relationship across eukaryotes

To evaluate the coupling between abundance and genetic diversity, we quantified three metrics for each OTU within each phylum: (i) relative abundance (log-transformed read counts, used as a proxy for abundance in metabarcoding studies (Lamb *et al*. 2019)), (ii) nucleotide diversity (π), and (iii) log-connections, a locality-level index derived from haplotype networks that captures the connectivity of each locality cluster within each phylum-specific network. We then computed pairwise Spearman correlations (ρ) between these metrics using raw values (Figure 2a). Across phyla, plants and animals exhibited consistently stronger correlations between abundance and nucleotide diversity than protists (mean ∼42% vs. ∼30%; Figure 2b, Table S1). This contrast became sharper when grouping taxa by cellularity: multicellular organisms showed substantially stronger coupling than unicellular ones (∼41% vs. ∼30%; Figure 2b, Table S2). Among unicellular groups, photosynthetic, the bloom-forming microalgal groups Chlorophyta (ρ = 0.26), Haptophyta (ρ = 0.30) and Myzozoa (ρ = 0.20) displayed particularly weak associations (Figure 2a).

**Figure 2.**
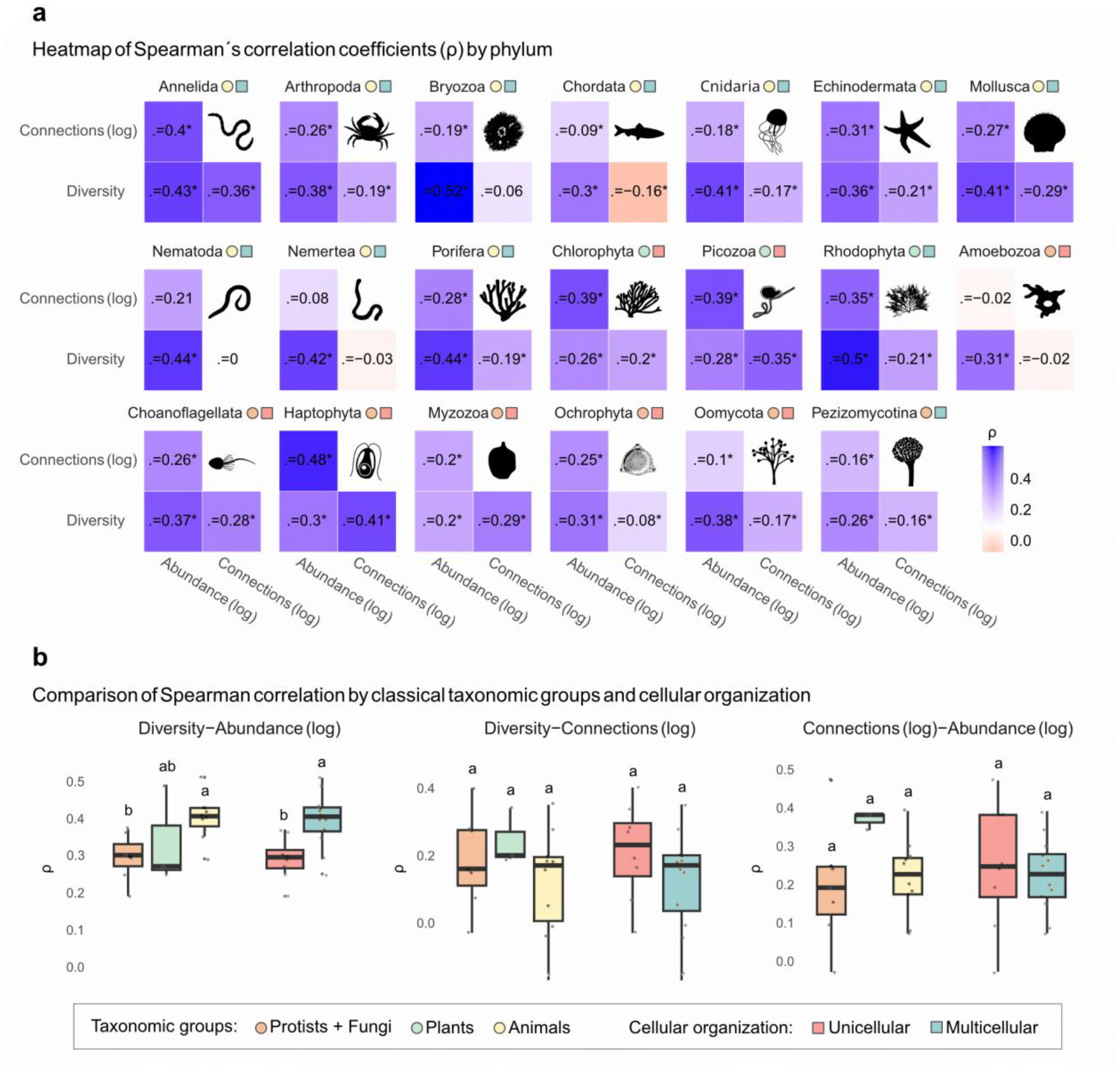
Relationships among abundance, nucleotide diversity and haplotype-network connectivity across marine eukaryotic phyla. **a**, Phylum-level heatmaps of pairwise Spearman correlation coefficients (ρ) among log-transformed relative abundance, nucleotide diversity and haplotype-network connectivity, expressed as log-connections. Cell colours represent correlation strength, and asterisks indicate statistically significant correlations. **b**, Group-level comparisons of Spearman correlation coefficients for diversity-abundance, diversity-connectivity and connectivity-abundance relationships across taxonomic groups and unicellular versus multicellular lineages. Boxes represent interquartile ranges, horizontal lines denote medians, whiskers show the data range excluding outliers, and points correspond to individual phyla. Different letters indicate significant differences among groups.

By contrast, correlations involving haplotype-network connectivity were weaker and less structured across cellularity types (Figure 2b, Table S2). Differences between multicellular and unicellular lineages were modest for both abundance vs. connections (∼23% vs. ∼28%) and connections vs. π (∼14% vs. ∼17%). However, some particular phyla exhibited relatively high connectivity correlations, including Haptophyta (abundance vs. haplotype connections 48%, and haplotype connections vs. π, 41%), Annelida (40% and 36%), Chlorophyta (39% and 20%), Picozoa (39% and 35%), and Rhodophyta (35% and 21%) (Fig. 2a).

### 3.3. Distance-decay patterns

We next assessed how abundance, nucleotide diversity, and network connectivity varied with geographic distance. Across phyla, exponential decay models (NLS) consistently provided the best fit (Figure S1). This pattern is consistent with recent expansion dynamics, in which diversity and connectivity are concentrated around a limited number of close core regions. Although recently colonized localities tend to exhibit lower genetic diversity and stronger differentiation due to serial founder effects and insufficient time for divergence-migration equilibrium to develop (González-Miguéns *et al*. 2026; Peter & Slatkin 2015).

We further compared distance-decay slopes among phyla using permutation tests. Relative abundance showed the strongest deviations from null expectations in the meiofaunal metazoan phyla (Bryozoa, Nematoda, and Nemertea), with weaker but still significant effects in the mostly benthic and multicellular algae (Rhodophyta) and metazoa (Echinodermata, Mollusca) (Figure S2). Nucleotide diversity displayed a largely similar pattern, with the strongest deviations observed in Bryozoa, Chordata, Echinodermata, Nemertea, Nematoda, and Rhodophyta (Figure S3). Network connectivity also followed this trend, with the largest departures from null expectations detected in Bryozoa, Nematoda, Pezizomycotina, and Rhodophyta (Figure S4). Across all metrics, meiofaunal metazoan phyla (Bryozoa, Nematoda, Nemertea) and mostly benthic multicellular algae (Rhodophyta) consistently exhibited the strongest deviations, suggesting that life-history traits may play a more important role than cellularity itself in shaping spatial genetic structure.

Within each phylum, comparisons of distance-decay slopes among metrics revealed frequent and significant differences (Figure S5). In most cases, abundance declined more steeply with geographic distance than nucleotide diversity or network connectivity. This decoupling was particularly pronounced among meiofaunal taxa, where abundance exhibited strong spatial decay, whereas genetic diversity and connectivity remained comparatively stable or heterogeneous.

Taken together, both correlation analyses and distance-decay models reveal a consistent but incomplete coupling between relative abundance and nucleotide diversity across eukaryotes. Although this relationship is generally positive, it is weaker in unicellular lineages. These results indicate that, in microbial eukaryotes, classical biodiversity metrics based on abundance or occurrence cannot be assumed to capture the evolutionary and demographic processes underlying diversity patterns. Therefore, our results highlight the importance of integrating population genetic information into ecological and biogeographic frameworks. Metrics such as genetic connectivity and intraspecific diversity provide complementary signals on dispersal, demographic structure, and evolutionary dynamics that are not captured by traditional community-level metrics.

### 3.4. Local connectivity networks

Because abundance-based patterns alone do not fully capture the evolutionary and demographic processes underlying diversity across eukaryotes, we next examined how intraspecific genetic diversity is spatially structured across marine systems. To do so, it is necessary to integrate information on both genetic relatedness and geographic distribution. Network-based approaches provide a powerful framework for representing these relationships, allowing spatial patterns of diversity to be quantified below the OTU level (González-Miguéns *et al*. 2026; Turon *et al*. 2020). Here, we developed a locality-centred approach that transforms haplotype networks into spatial connectivity networks (Figure 1b) (see Materials and Methods and Figure S6).

To investigate whether haplotype-network connectivity captures biologically meaningful spatial structure across marine biogeographic transitions, an ideal system should encompass well-defined oceanographic boundaries and contrasting environmental regions. In this regard, the Iberian Peninsula provides an excellent natural setting, as it includes a sharp transition between the temperate North Atlantic Ocean and the subtropical Mediterranean Sea (Figure 3a). Using this well-characterized biogeographic region, we compared the geographic distances associated with haplotype-network connections across taxonomic and cellularity-based categories. Under a scenario of spatial genetic structuring, connections linking distinct haplotypes (“different”) are expected to span larger geographic distances than those linking identical haplotypes (“same”), reflecting increasing genetic differentiation with geographic separation (Figure S7). Therefore, we quantified the median geographic distance associated with each type of pairwise haplotype connection, allowing genetic similarity and spatial structure to be integrated within a unified analytical framework.

**Figure 3.**
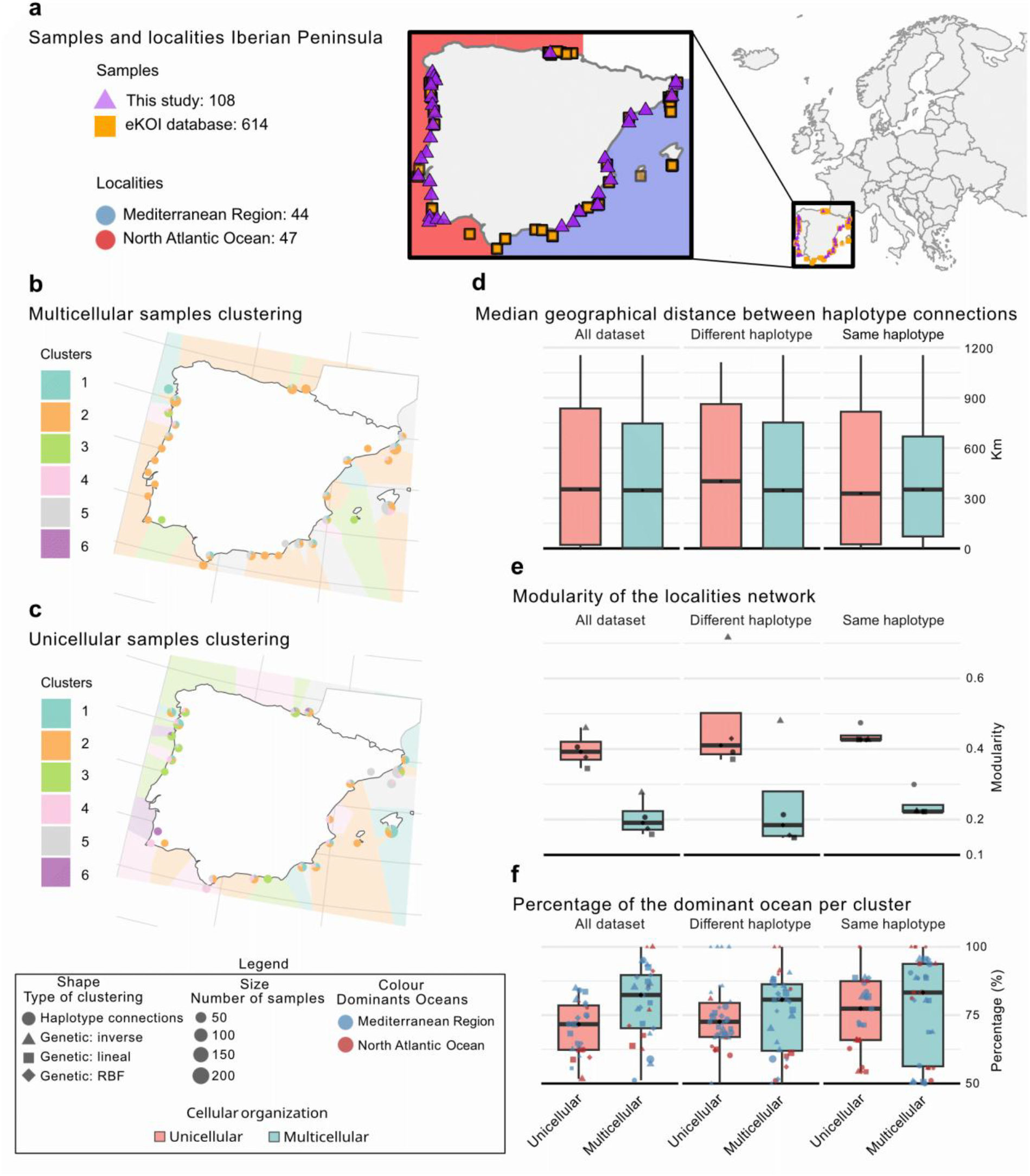
Geographic haplotype networks from metabarcoding data. **a**, Samples from the Iberian Peninsula obtained from the eKOI metabarcoding database 2.0 (squares) and newly generated in this study (triangles). **b**, Clustering of Iberian localities using haplotype-network connections for multicellular organisms, and **c**, equivalent clustering for unicellular organisms. **d**, Boxplots showing the median geographic distance between all connections (“all”), only intra-haplotype connections (“same”), and inter-haplotype connections (“different”) for unicellular and multicellular groups. **e**, Boxplots representing the modularity values of the locality networks for each type of connection and cell-type category. **f**, Boxplots showing the percentage of the dominant ocean (Mediterranean or North Atlantic) within each locality cluster, separated by connection type and cell-type category.

Across the Iberian Peninsula dataset, differences among taxonomic groups (Figure S8, Data S4) were generally modest. Plants showed slightly shorter median connection distances (median =289 km) than protists (347 km) and animals (353 km). Comparisons between “same” and “different” classes indicated that “different” connections tended to span larger geographic distances in protists (−93 km difference), whereas contrasts were weak or inconsistent in both Plants (+11 km) and Animals (+38 km). Overall, these differences were small, suggesting broadly comparable spatial structuring across eukaryotes.

Differences were even smaller when taxa were grouped by cellularity (Figure 3, Data S5). Across the Iberian Peninsula dataset, unicellular organisms (median = 353 km) were only slightly higher than multicellular ones (347 km). The “same” “different” difference was again consistent but less marked: Unicellular (−73 km) > Multicellular (+5 km). Taken together, these results indicate that connections among distinct haplotypes (“different”) tend to occur across larger geographic distances, particularly in protists and unicellular taxa, consistent with a slightly weaker pattern of isolation by distance at the haplotype-network level. However, the limited magnitude of these differences suggests broadly similar biogeographic structuring across unicellular and multicellular eukaryotes.

### 3.5. Locality network cluster structure and modularity

We next inquired whether intraspecific genetic connectivity generates coherent spatial organization among localities. To do this, we transformed haplotype-network connections into locality-centred graphs. In these networks, localities are connected according to the genetic proximity among the haplotypes they contain, allowing the spatial structure of genetic diversity to be represented as interconnected geographic networks. This framework enables both the identification of clusters of highly connected localities and the quantification of large-scale network organization.

To characterize these patterns, we identified clusters of interconnected localities using the Louvain community detection algorithm and quantified network modularity across alternative edge-weighting approaches (Blondel *et al*. 2008; Newman 2004). We evaluated four weighting schemes: (i) connection count (unweighted genetic intensity), (ii) inverse genetic distance, (iii) linear genetic weighting, and (iv) radial basis function (RBF) weighting (Figures S9 and S10).

Across all weighting schemes, protists/fungi consistently exhibited higher modularity (median > 0.4) than plants and animals (median < 0.3) (Figure S8, Table S3). This pattern became even more pronounced when taxa were grouped by cellularity (Fig. 3, Table S3), with unicellular lineages showing substantially higher modularity (∼0.4) than multicellular ones (∼0.2). These results indicate stronger spatial compartmentalization and internal genetic cohesion in unicellular lineages, despite their often-weaker coupling between abundance and nucleotide diversity.

To assess whether these network modules also displayed geographic coherence, we calculated Moran’s I (Figure S11, Table S4) for cluster membership (k = 5 nearest neighbours). All clustering schemes showed positive spatial autocorrelation, with values typically between 0 and 0.3, indicating moderate spatial structure. Within the taxonomic groups, animals displayed the highest median Moran’s I across all clustering schemes (all = 0.192, same = 0.098, different = 0.126), followed by plants (0.147, 0.073, 0.126) and protists (0.095, 0.165, ≈0.000). Differences between cellularity groups were comparatively small: unicellular lineages showed Moran’
ss I value of 0.188 (all), 0.308 (same), and 0.020 (different), whereas multicellular lineages showed 0.192, 0.234, and 0.120, respectively. As expected, clusters derived from “different” connections tended to show lower spatial coherence than those derived from “same” connections, consistent with the idea that genetically distinct haplotypes connect more distant localities. Overall, differences between unicellular and multicellular lineages remained modest, suggesting broadly comparable levels of spatial aggregation.

### 3.6. Biogeographic partitioning

Finally, as a consistency check, we tested whether the network-derived locality clusters reflected known biogeographic boundaries between major oceanic regions. If locality-centred connectivity networks capture biologically meaningful patterns of population structure, clusters would be expected to align with well-documented marine biogeographic regions rather than forming geographically mixed assemblages. Consistent with this expectation, most locality clusters were strongly dominated by a single oceanic region (>70% of localities), indicating high geographic coherence (Figure S12, Table S4). Animals showed ocean-dominated clusters in 81.2% (all haplotypes), 81.2% (same haplotypes), and 78.3% (different haplotypes) of cases; plants in 75.1%, 63.9%, and 85.7%; and protists in 77.9%, 68.3%, and 71.4%, respectively (Figure S8). Under the cellularity grouping, multicellular lineages showed more ocean-homogeneous clusters (85.5%, 83.3%, 81.2%) than unicellular ones (74.3%, 71.4%, 74.4%), indicating clearer large-scale oceanic partitioning (Figure 3f). In contrast, unicellular lineages more frequently formed clusters spanning multiple marine regions. Across both grouping schemes, clusters were more frequently dominated by the Mediterranean region than by the North Atlantic (animals: Mediterranean 21 vs. Atlantic 8 localities clusters; plants: 16 vs. 8; protists: 18 vs. 8; multicellular: 20 vs. 9; unicellular: 18 vs. 7). This pattern suggests that the Mediterranean coast harbours greater intraspecific genetic diversity than the northern Atlantic margin of the Iberian Peninsula across both unicellular and multicellular organisms.

Together, these results reveal contrasting but complementary patterns across eukaryotes. Unicellular taxa exhibit stronger internal network cohesion and modularity, whereas multicellular organisms show clearer large-scale biogeographic partitioning. However, the differences observed are moderate, and both groups display broadly similar spatial organization of genetic diversity across marine systems. More broadly, this framework provides a scalable approach to infer population connectivity and biogeographic structure, taking advantage of the high amount of data obtained from metabarcoding and its universality. This framework will help bridge the gap between community ecology and population genetics.

## 4. Discussion

Our results reveal a consistent but incomplete coupling between relative abundance and genetic diversity across marine eukaryotes. While a positive relationship is generally observed, it is markedly weaker in unicellular lineages than in multicellular organisms. These findings suggest that relative abundance, at least when approximated from metabarcoding reads, does not reliably capture population genetic processes in microbial eukaryotes.

This decoupling likely reflects fundamental differences in life-history traits. Many protists are characterized by rapid population growth and high dispersal potential, which can produce large and transient fluctuations in local abundance without corresponding changes in long-term effective population size. Bloom-forming taxa provide a clear example: groups such as Chlorophyta, Haptophyta, and Myzozoa frequently reach extremely high local abundances, yet show relatively weak associations between abundance and nucleotide diversity (Filatov 2019; Krueger-Hadfield *et al*. 2014; Rengefors *et al*. 2017). In the early stages of blooms, extremely high division rates lead to homogeneous populations, although genetic diversity increases later, driven by selective sweeps (Ryderheim & Kiørboe 2024).

At the same time, all three metrics analysed (abundance, nucleotide diversity, and haplotype-network connectivity) show a clear exponential decay with geographic distance. This pattern is consistent with recent or recurrent expansion dynamics in marine systems, in which diversity and connectivity are concentrated around a limited number of source regions. While anthropogenic transport may contribute to long-distance dispersal (González-Miguéns *et al*. 2026), similar patterns can also arise from natural oceanographic processes (Cowen *et al*. 2015; Selkoe *et al*. 2016) and post-glacial expansions (Maggs *et al*. 2008). In either case, the resulting genetic structure is shallow relative to geographic distance, reinforcing the observed decoupling between spatial distribution and genetic differentiation.

Together, these results challenge the widespread assumption that sequence abundance directly reflects ecological success or population stability. Here, our analyses show that highly abundant OTUs are not necessarily more genetically diverse, particularly in unicellular taxa. In contrast, intraspecific genetic diversity, shaped by mutation, drift, and gene flow, provides a more direct window into population stability and historical connectivity. Abundance and genetic diversity, therefore, capture partially independent dimensions of biodiversity and should not be used interchangeably.

A key contribution of this study is the introduction of a haplotype network-based framework that links intraspecific genetic variation to geographic space. By transforming haplotype relationships into locality-level connectivity networks, this approach enables the joint analysis of genetic similarity, spatial structure, and dispersal processes. Previous studies have demonstrated population structuring in metabarcoding data at specific taxonomic or geographic scales (González-Miguéns *et al*. 2026; Thomasdotter *et al*. 2023; Turon *et al*. 2020); here we extend this perspective across multiple phyla and spatial scales, providing a generalizable framework for inferring population-level processes from environmental sequencing data.

Applying this framework to the Iberian Peninsula reveals clear biogeographic patterns. Most locality clusters are dominated by a single oceanic region, indicating strong large-scale spatial structuring, particularly in multicellular taxa. At the same time, unicellular lineages exhibit higher network modularity, suggesting stronger internal cohesion and localized structuring despite their broader spatial spread. These results highlight a key contrast: multicellular organisms show clearer geographic partitioning, whereas unicellular taxa exhibit more cohesive but spatially diffuse population structures, perhaps due to long-range dispersal events that are not necessarily successful (Cowen & Sponaugle 2009; Martiny *et al*. 2006).

Despite these differences, both unicellular and multicellular groups share broadly similar spatial patterns in the distribution of genetic diversity, connectivity, and spatial autocorrelation. This convergence suggests that common biogeographic processes operate across all eukaryotes, challenging the traditional dichotomy between “microbial” and “macrobial” systems. Instead, the strength and spatial expression of these patterns seem to depend more strongly on lineage-specific life-history traits, such as dispersal mode, reproductive strategy, or habitat association, than on cellularity itself. Rather than representing fundamentally distinct biogeographic systems, unicellular and multicellular organisms may therefore occupy different positions along a shared continuum of spatial genetic organization. This is particularly evident in meiofaunal metazoans (e.g., Nematoda, Bryozoa, Nemertea), which consistently deviate from broader patterns and display strong spatial structuring across all metrics.

Although our approach represents a novel and scalable framework for integrating population genetic information into ecological and biogeographic analyses, it also entails several limitations that need to be considered. First, it relies on a single mitochondrial marker, which may introduce lineage-specific biases and does not capture the full complexity of genome-wide variation. Second, sequence-derived relative abundance cannot be directly interpreted as effective population size, as metabarcoding abundances are influenced by multiple technical and biological factors, including biomass, copy-number variation, and amplification biases. Nevertheless, relative abundance remains an informative ecological proxy for broad-scale comparisons of dominant and rare lineages across marine systems. Third, haplotype-network reconstruction depends on methodological choices that can influence inferred connectivity patterns. Finally, although we focus on cellularity as a broad organizing principle, other ecological and evolutionary traits are likely to be equally or more important in shaping spatial genetic structure.

Despite these limitations, our results demonstrate that integrating haplotype-level information with spatial data provides a powerful avenue to move beyond community composition and towards a process-based understanding of biodiversity. By explicitly linking genetic variation to geographic structure, our framework bridges “metabarcoding-based” ecology and population genetics, enabling the inference of dispersal, connectivity, and demographic history across diverse lineages.

More broadly, this work contributes to a growing effort to extend population-genetic principles to microbial eukaryotes. While classical expectations linking abundance and genetic diversity are not fully sustained, they are not absent either. Instead, they are modulated by life-history traits, spatial dynamics, and temporal scales. Recognizing and quantifying this partial decoupling is essential for developing a unified, cross-lineage theory of biodiversity that encompasses both unicellular and multicellular life.

## Acknowledgements

We thank Emilio Cano Cabezas for his help at the lab; Alejandro Berlinches de Gea, Carmen Soler-Zamora, Fernando Useros, Nura ElKhouri Vidarte and Ángel García Bodelón for fruitful discussions. We are also grateful to all members of the Multicellgenome Lab (MCG) for their assistance and for fostering such a supportive working environment.

## Statement of authorship

Conceptualisation: M.M.-R., I.R.-T., E.L. and R.G.-M. Methodology, software, and formal analysis: M.M.-R., P.G.P and R.G.-M Validation: M.M.-R, P.G.P and E.L. Investigation: M.M.-R., I.R.-T, E.L. and R.G.-M Data curation: M.M.-R., P.G.P., and R.G.-M Project administration and funding acquisition: I.R.-T. and E.L. Writing - original draft: M.M.-R and R.G.-M. Writing - review and editing: All authors.

## Data accessibility statement

Processed datasets, supplementary figures and supplementary tables are available from Zenodo at DOI: 10.5281/zenodo.21392170. Raw COI metabarcoding sequence data generated for this study have been deposited in the NCBI Sequence Read Archive under BioProject accession PRJNA1493043. Further implementation details and the complete code used in the analyses are available in the GitHub repository pablogpena/phylaCOI. The repository includes modular command-line scripts and a Nextflow (Di Tommaso et al. 2017) implementation of the full analysis, enabling reproducible execution across computing environments.

