## Supplementary Information for "Widespread decoupling between abundance and genetic diversity, and strong local genetic structuring in marine unicellular eukaryotes"

**Supplementary Files**

Data availability note. Supplementary Tables S1-S4 and Supplementary Data S1-S5 are deposited in Zenodo and are publicly available at https://doi.org/10.5281/zenodo.21392170. Their captions and file descriptions are provided below.

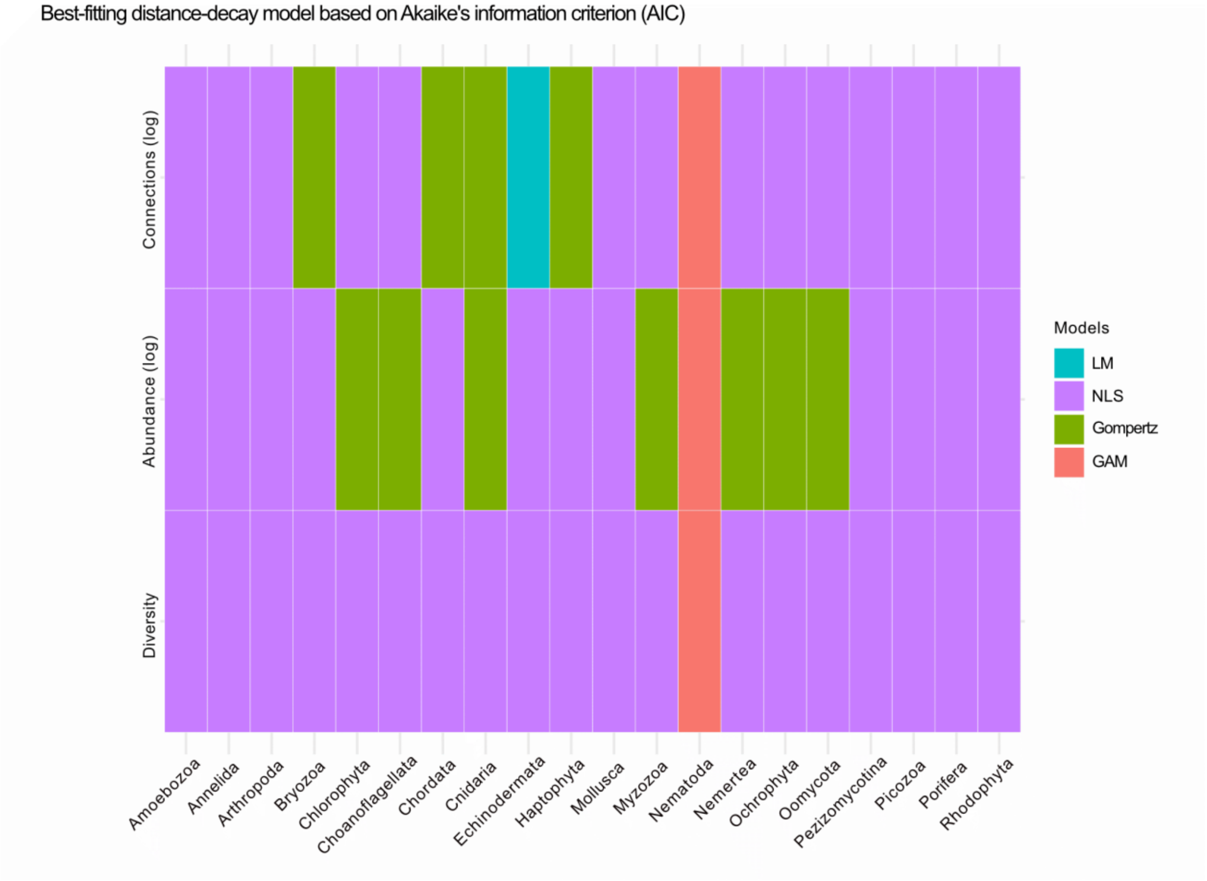


**Figure S1. Best-fitting distance-decay model for each phylum and metric, selected using Akaike's information criterion (AIC)**. Rows show log-transformed haplotype-network connectivity, log-transformed relative abundance and nucleotide diversity, and columns show eukaryotic phyla. Colours indicate the selected model: linear model (LM), nonlinear least-squares model (NLS), Gompertz model or generalized additive model (GAM).

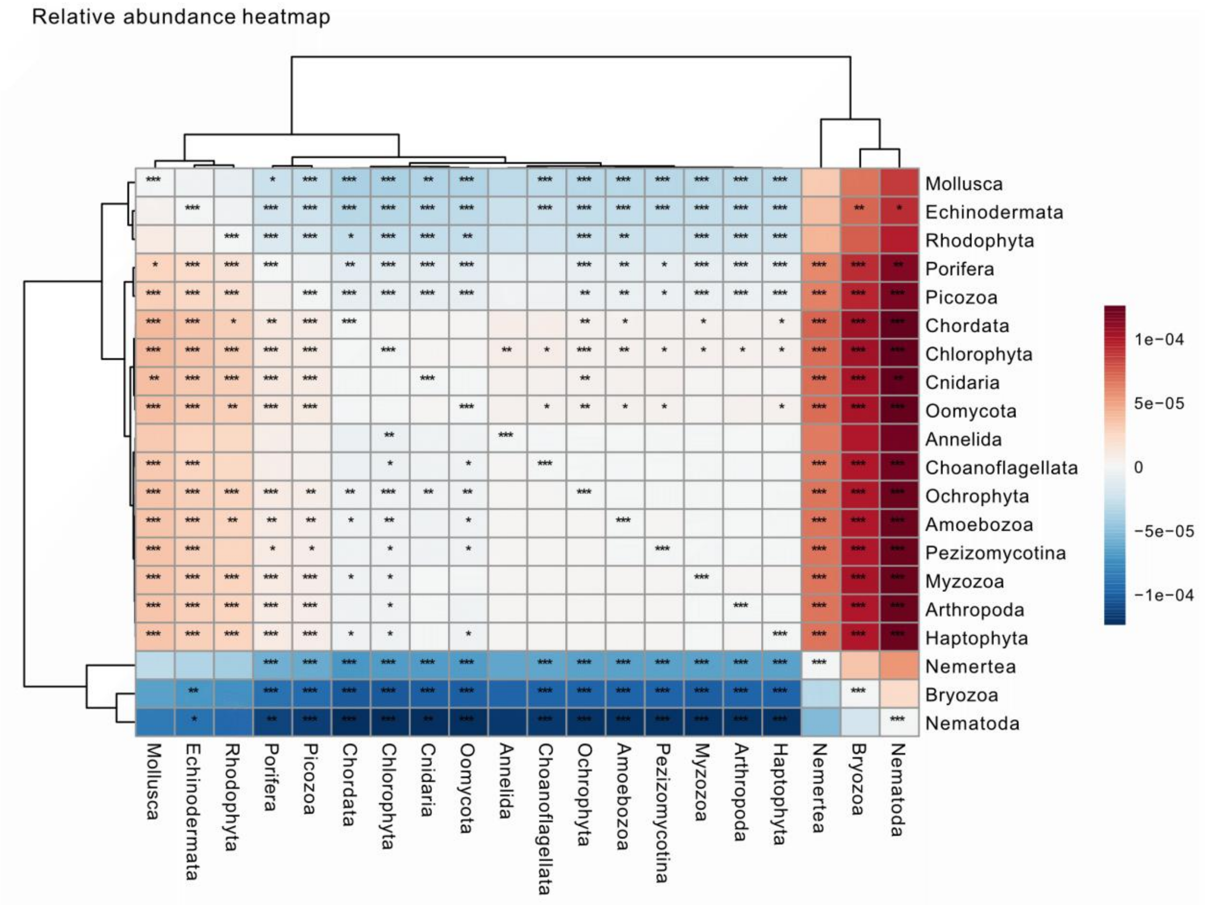


**Figure S2. Heatmap showing pairwise comparisons of distance-decay slopes for log-transformed relative abundance among eukaryotic phyla**. Colours indicate the direction and magnitude of pairwise slope differences, with rows and columns ordered by hierarchical clustering. Asterisks denote significant differences based on permutation tests after Benjamini-Hochberg false discovery rate correction ($p < 0.05$=*; $p < 0.01$=**; $p < 0.001$=***).

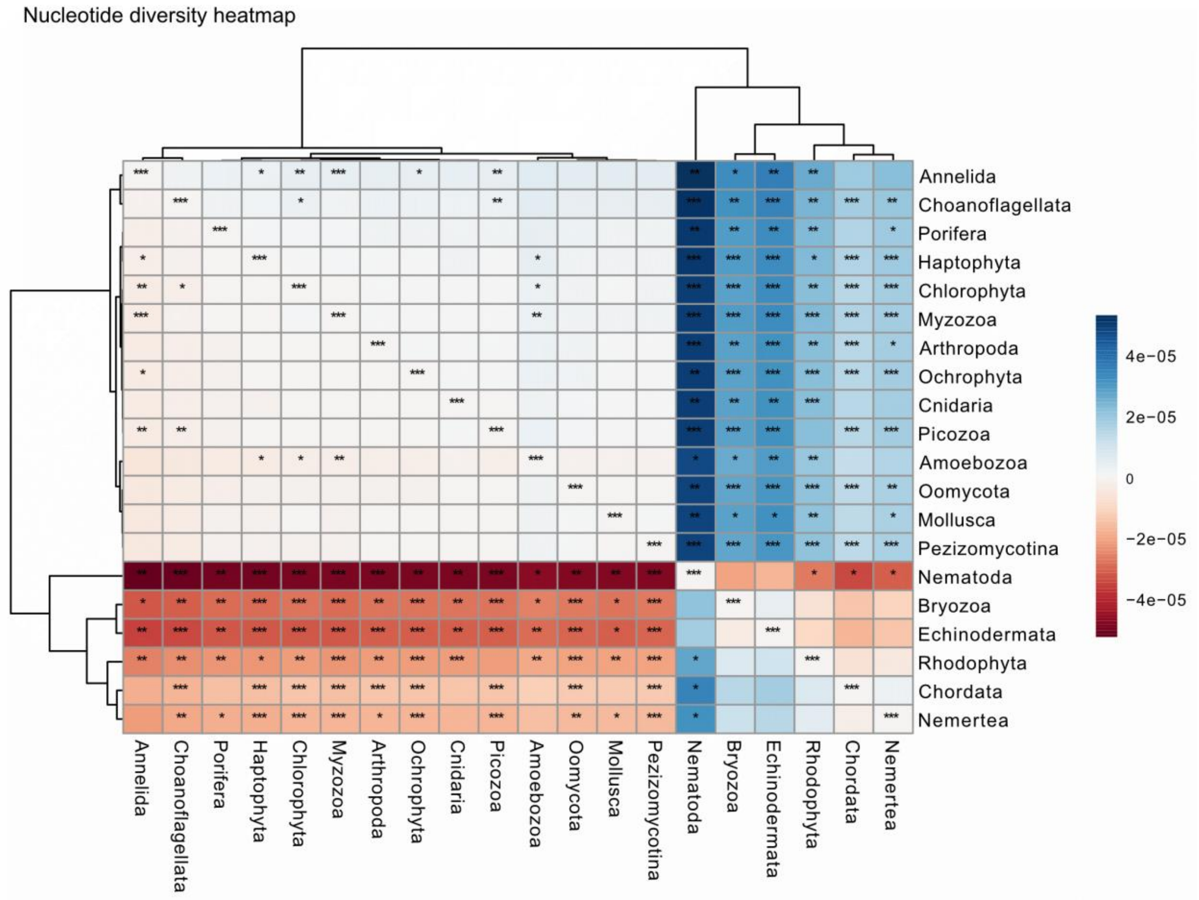


**Figure S3. Heatmap showing pairwise comparisons of distance-decay slopes for nucleotide diversity among eukaryotic phyla**. Colours indicate the direction and magnitude of the pairwise slope differences between phyla, with rows and columns ordered by hierarchical clustering. Asterisks denote significant differences based on permutation tests after Benjamini-Hochberg false discovery rate correction ($p < 0.05$=*; $p < 0.01$=**; $p < 0.001$=***).

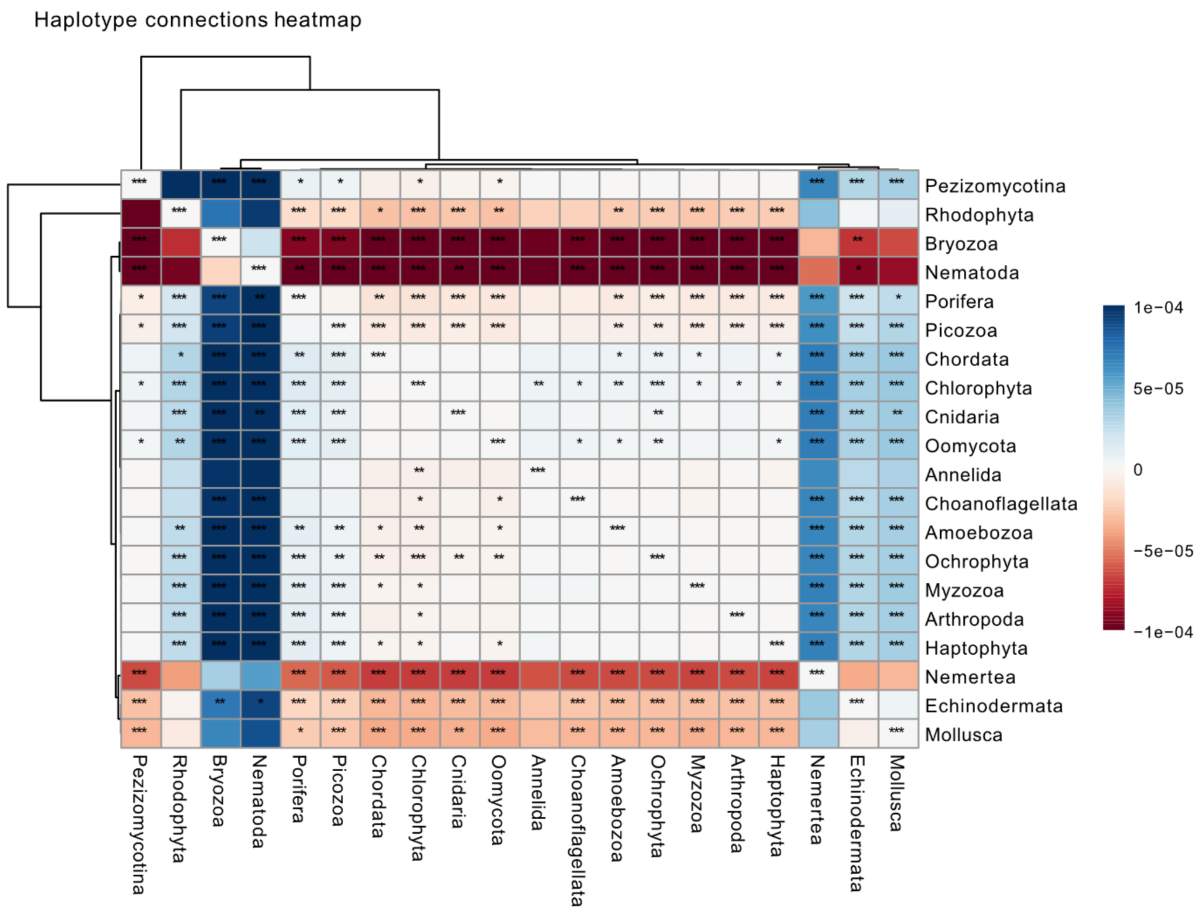


**Figure S4. Heatmap showing pairwise comparisons of distance-decay slopes for haplotype-network connectivity, measured as log-connections, among eukaryotic phyla**. Colours indicate the direction and magnitude of the pairwise slope differences between phyla, with rows and columns ordered by hierarchical clustering. Asterisks denote significant differences based on permutation tests after Benjamini-Hochberg false discovery rate correction ($p < 0.05$=*; $p < 0.01$=**; $p < 0.001$=***).

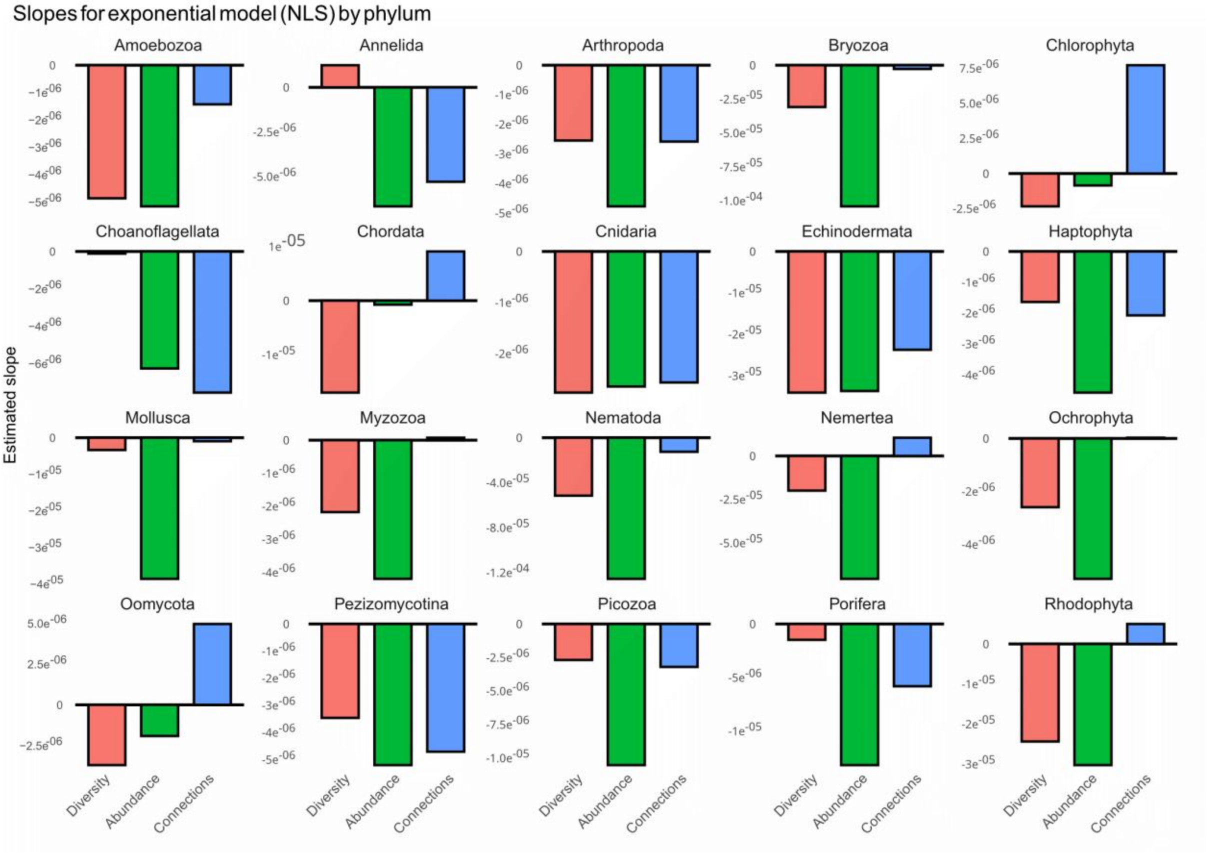


**Figure S5. Estimated distance-decay slopes from the exponential nonlinear least-squares model (NLS) for each eukaryotic phylum and metric**. Bars show the fitted slope values for nucleotide diversity (diversity), log-transformed relative abundance (abundance), and log-transformed haplotype-network connectivity (connections) measured as log-connections. Negative slopes indicate decreasing values with increasing geographic distance from the metric-specific centre, whereas positive slopes indicate increasing values with distance.

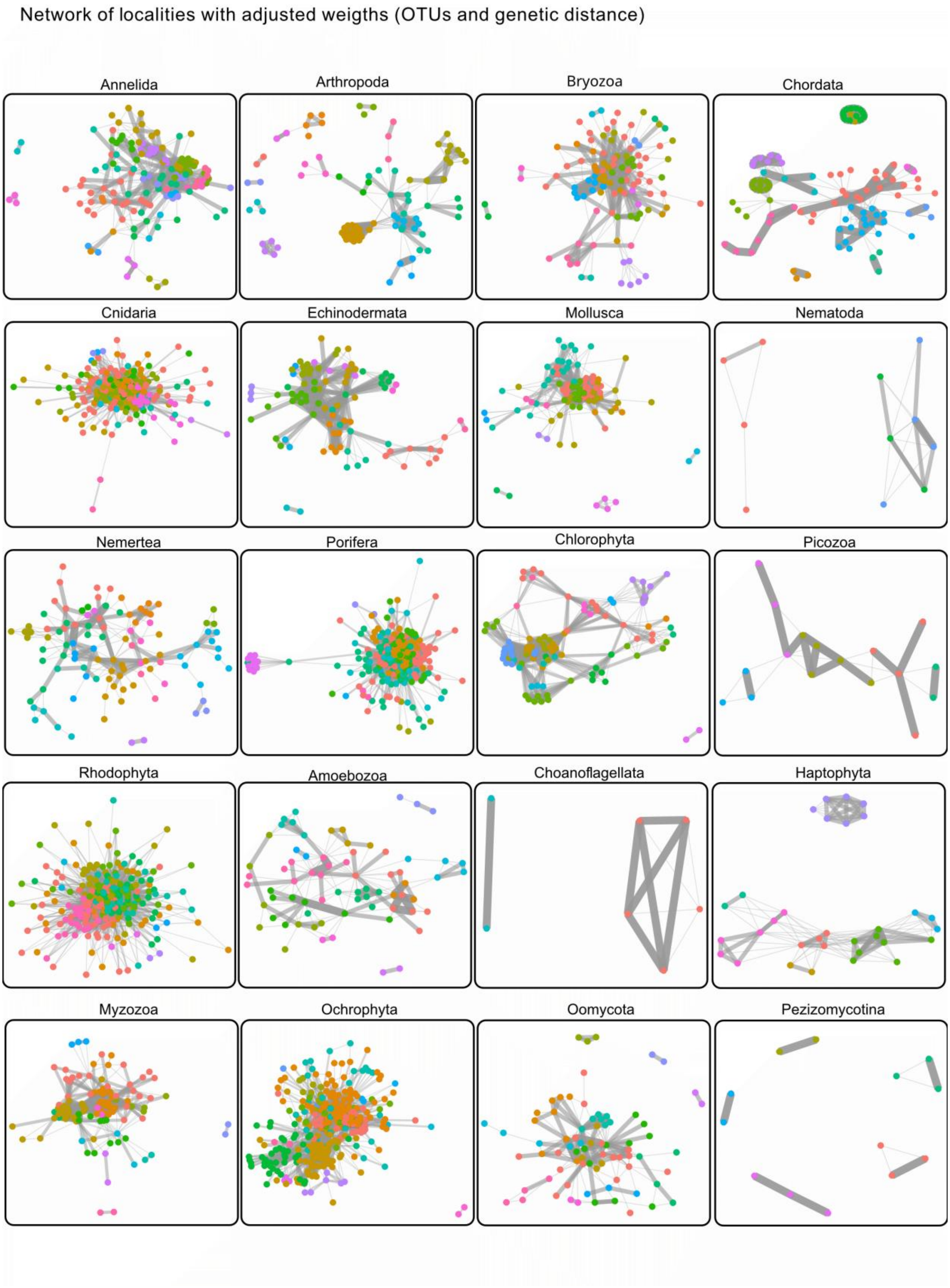


**Figure S6. Locality networks weighted by inverse genetic distance**. Networks of localities constructed for each phylum, where nodes represent localities and edges represent genetic connections between them. Edge thickness is proportional to the connection weight, calculated as the inverse of mean genetic distance between connected haplotypes.

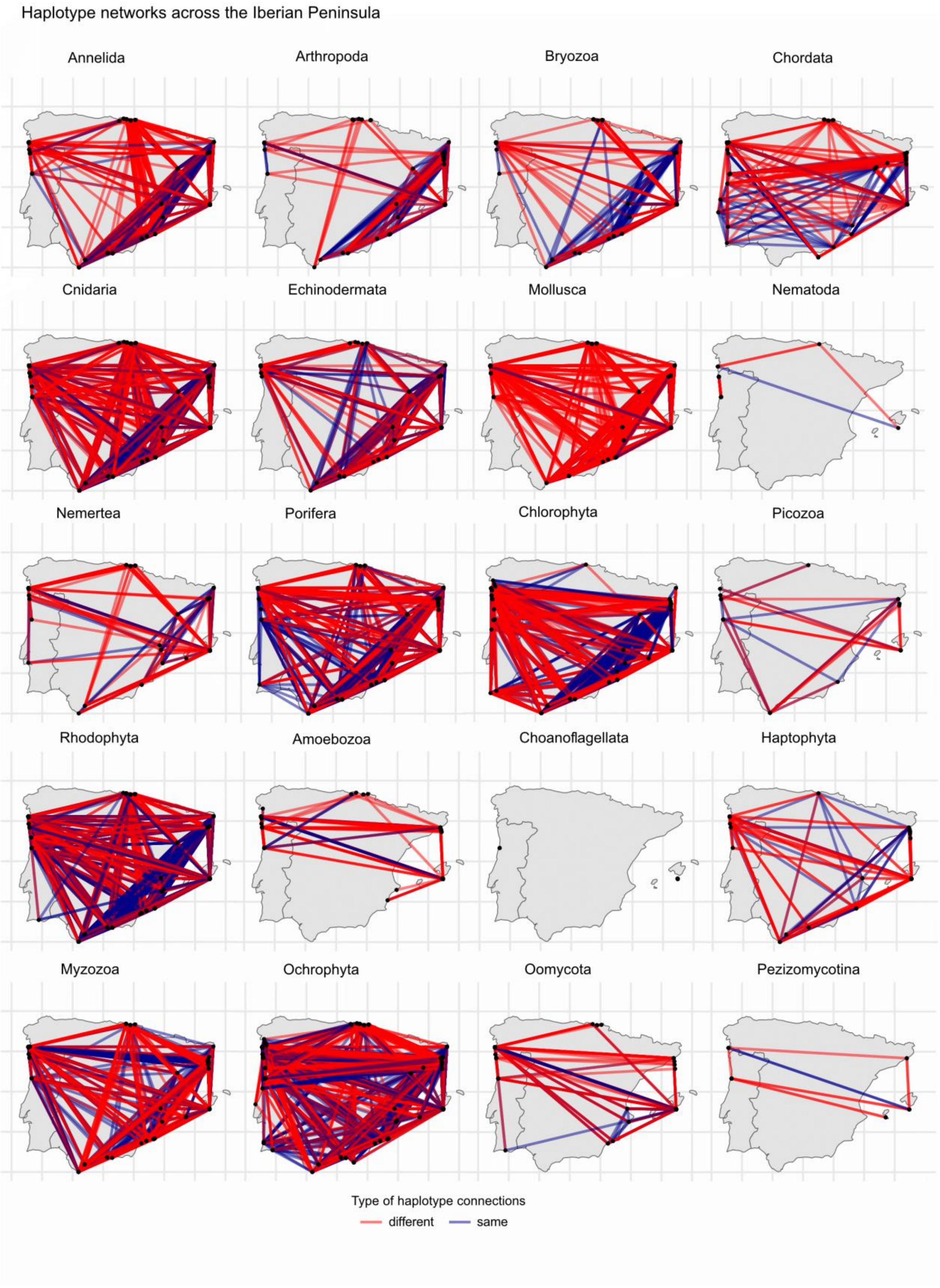


**Figure S7. Haplotype networks across the Iberian Peninsula combining OTUs by phylum**. Haplotype networks were represented for each phylum, showing the geographic distribution of all haplotype connections across the Iberian Peninsula: connections between different haplotypes ("different") in red, and connections within the same haplotype ("same") in blue.

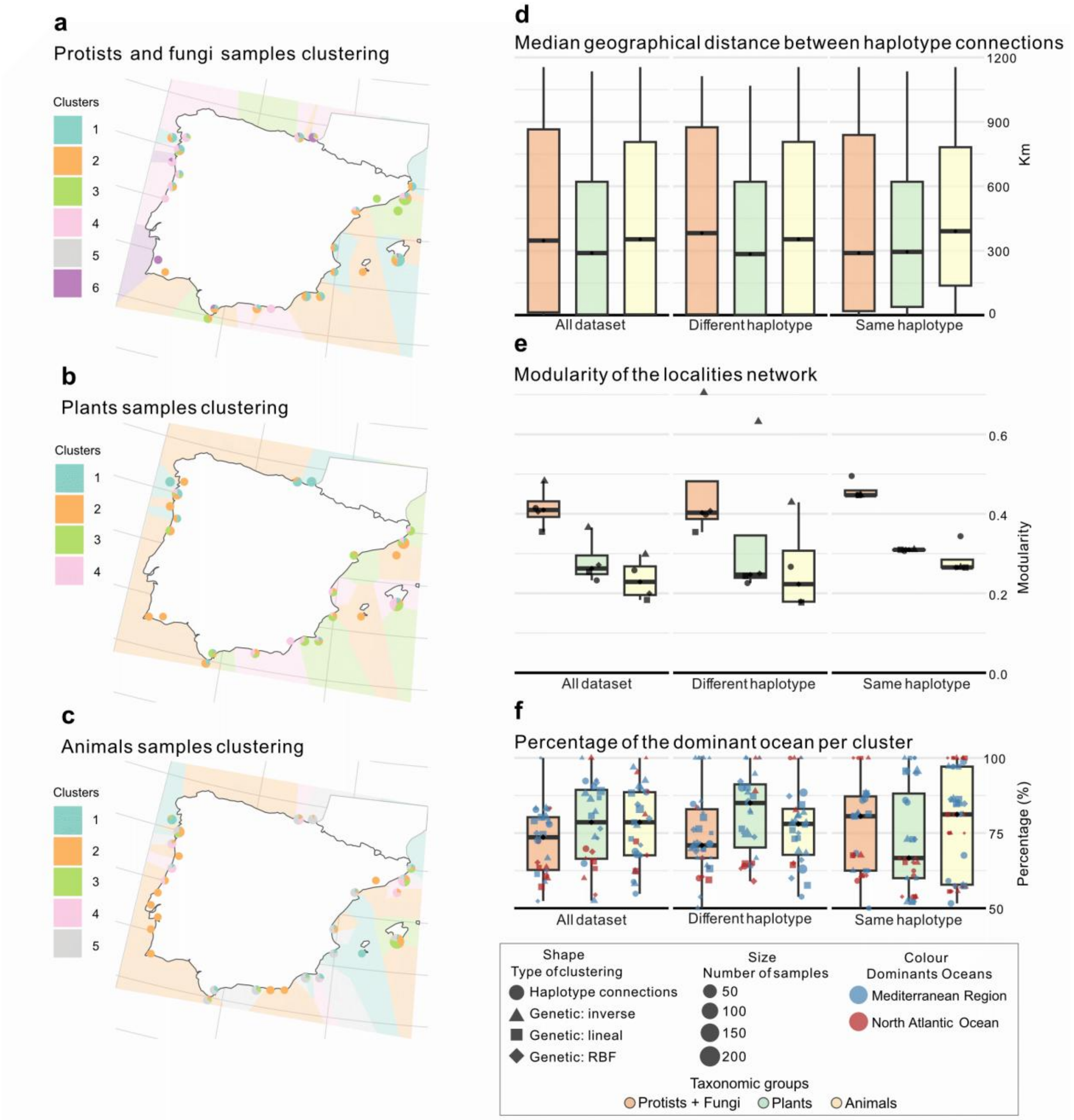


**Figure S8. Geographic haplotype networks by taxonomic groups**. **a,** Clustering of Iberian localities using haplotype-network connections for protist/fungi taxa; **b,** equivalent clustering for plant taxa; and **c,** equivalent clustering for animal taxa. **d,** Boxplots showing the median geographic distance between all connections ("all"), only intra-haplotype connections ("same"), and inter-haplotype connections ("different") for plants, animals and protists & fungi taxa. **e,** Boxplots representing the modularity values of the locality networks for each type of connection and taxonomy category. **f,** Boxplots showing the percentage of the dominant ocean (Mediterranean or North Atlantic) within each locality cluster, separated by connection type and taxonomy category.

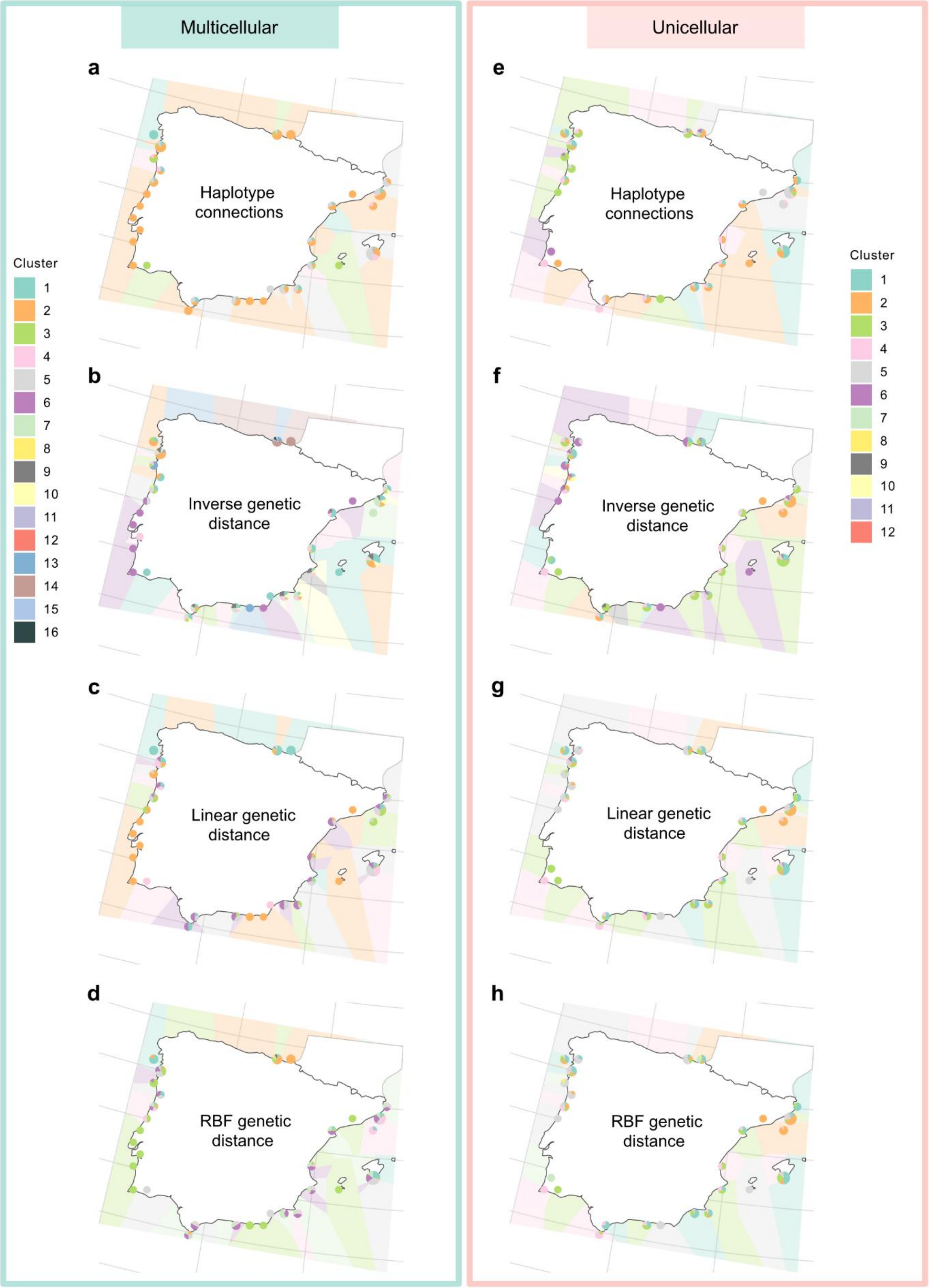


**Figure S9. Clustering of Iberian Peninsula localities by cell type**. **a-d**, Clustering of multicellular organisms based on haplotype-network information. Localities within a 5 km radius were grouped and color-coded according to their assigned cluster. Each locality is represented as a pie chart indicating the proportion of sites belonging to each cluster: **a,** clustering using haplotype-network connections only;

**b,** clustering weighted by the inverse of genetic distance; **c,** clustering weighted linearly by genetic distance; and **d,** clustering weighted by an RBF (radial basis function) transformation of genetic distance. **e-h**, Equivalent clustering for unicellular organisms, applying the same four methods: **e,** connections only; **f,** inverse genetic distance; **g,** linear genetic distance; and **h,** RBF-weighted genetic distance.

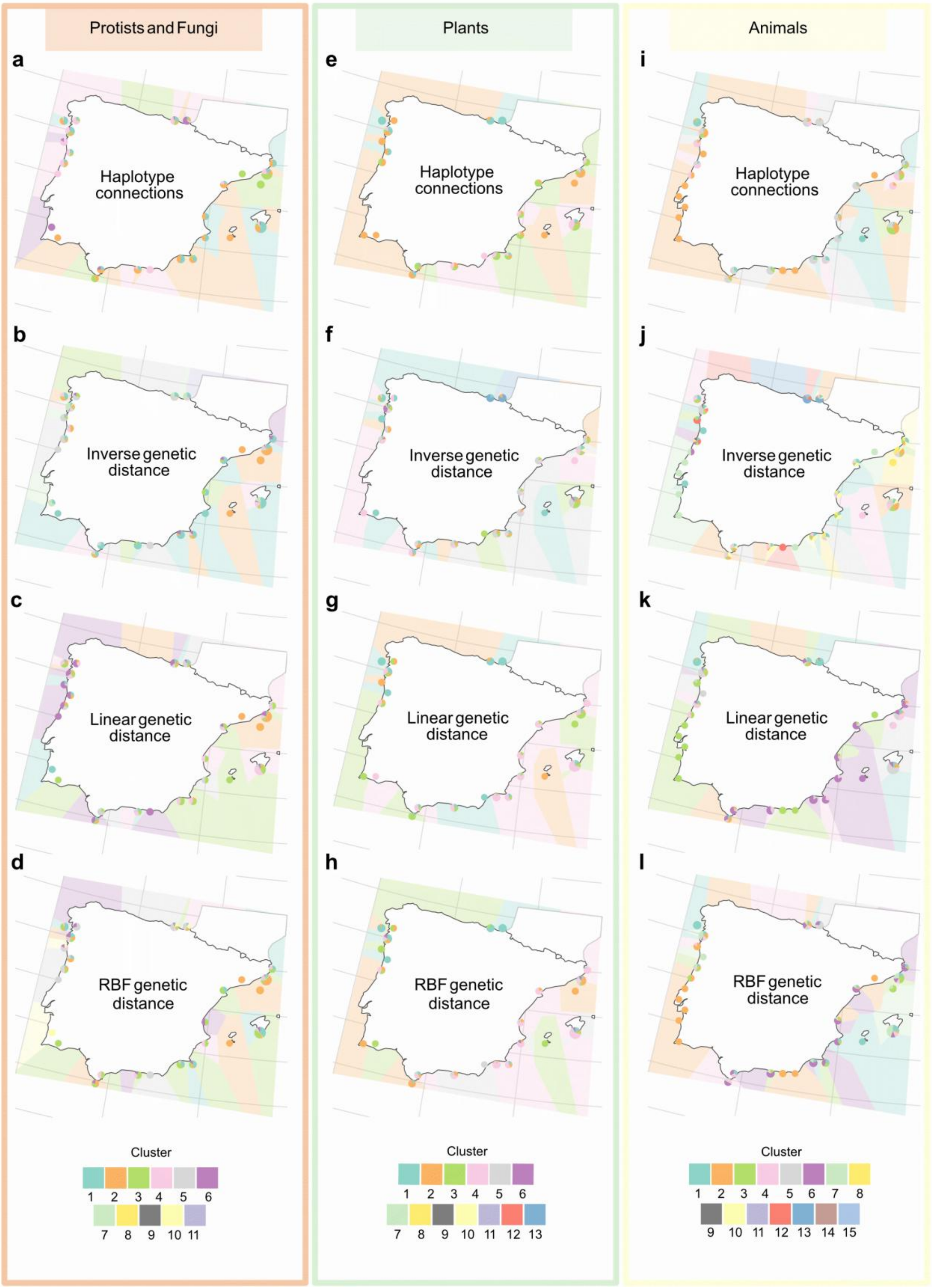


**Figure S10**. **Clustering of Iberian Peninsula localities by taxonomic groups**. **a-d**, Clustering of protist and fungi taxa based on haplotype-network information. Localities within a 5 km radius were grouped and color-coded according to their assigned cluster. Each locality is represented as a pie chart indicating the proportion of sites belonging to each cluster: **a,** clustering using haplotype-network connections only; **b,** clustering weighted by the inverse of genetic distance; **c,** clustering weighted linearly by genetic distance; and **d,** clustering weighted by an RBF (radial basis function) transformation of genetic distance. **e-h,** Equivalent clustering for plant taxa, applying the same four methods: **e,** connections only; **f,** inverse genetic distance; **g,** linear genetic distance; and **h,** RBF-weighted genetic distance. **i-l,** Equivalent clustering for animal taxa, applying the same four methods: **i,** connections only; **j,** inverse genetic distance; **k,** linear genetic distance; and **l,** RBF-weighted genetic distance.

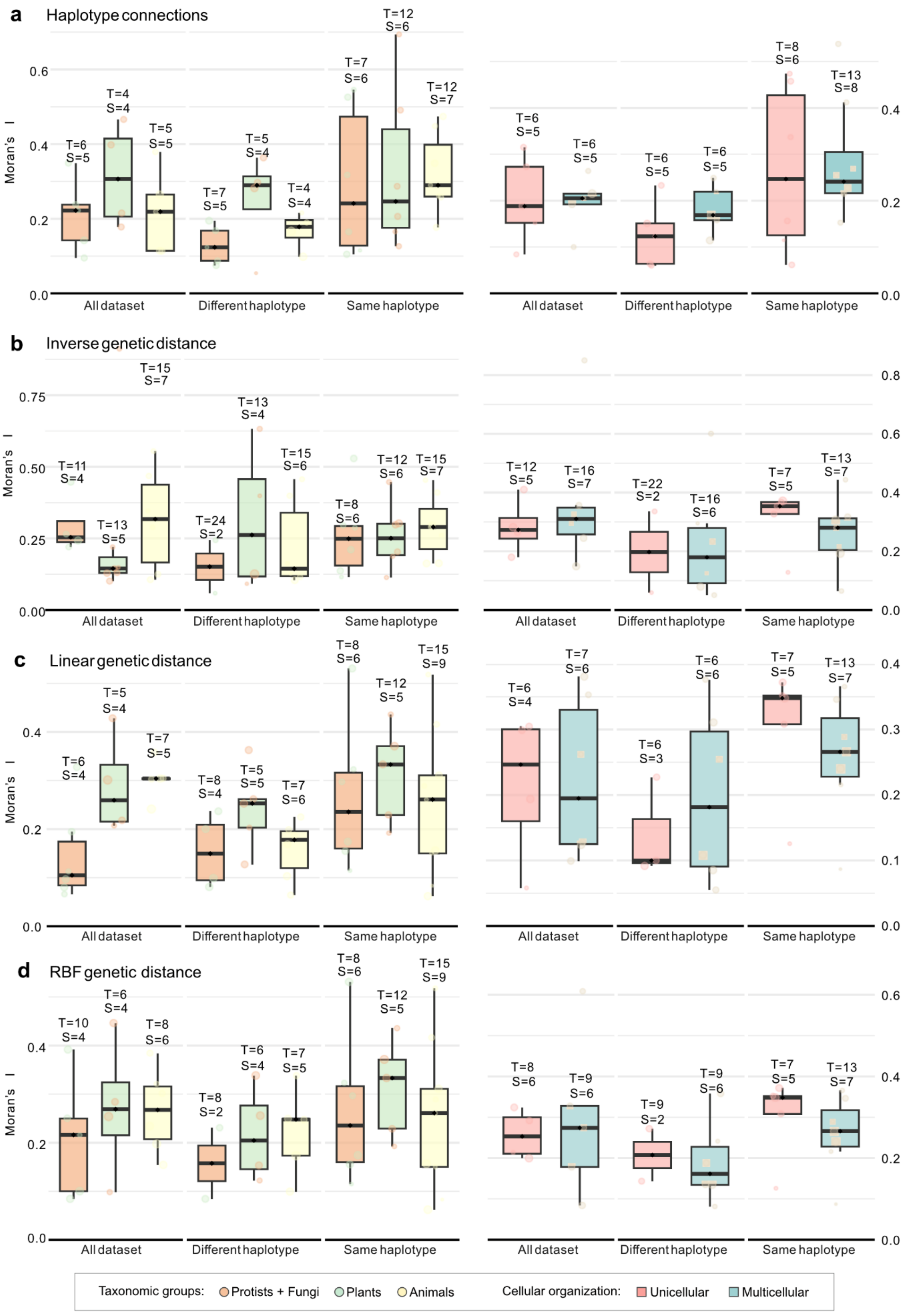
a
Haplotype connections
b
Inverse genetic distance
c
Linear genetic distance
d
RBF genetic distance
Moran's I
All dataset
Different haplotype
Same haplotype
Taxonomic groups: Protists + Fungi Plants Animals
Cellular organization: Unicellular Multicellular

**Figure S11**. **Spatial autocorrelation (Moran's I) of locality clusters**. Boxplots showing the Moran's I spatial autocorrelation index for locality clusters that are statistically significant and composed of at least three samples. Within each panel, results for taxonomic groups (protists and fungi, plants and animals) are shown on the left, whereas results according to cellular organization (unicellular and multicellular taxa) are shown on the right. Results are presented for networks constructed using: **a**, haplotype-network connections only; **b**, inverse genetic-distance weighting; **c**, linear genetic-distance weighting; and **d**, an RBF (radial basis function) transformation of genetic distance.

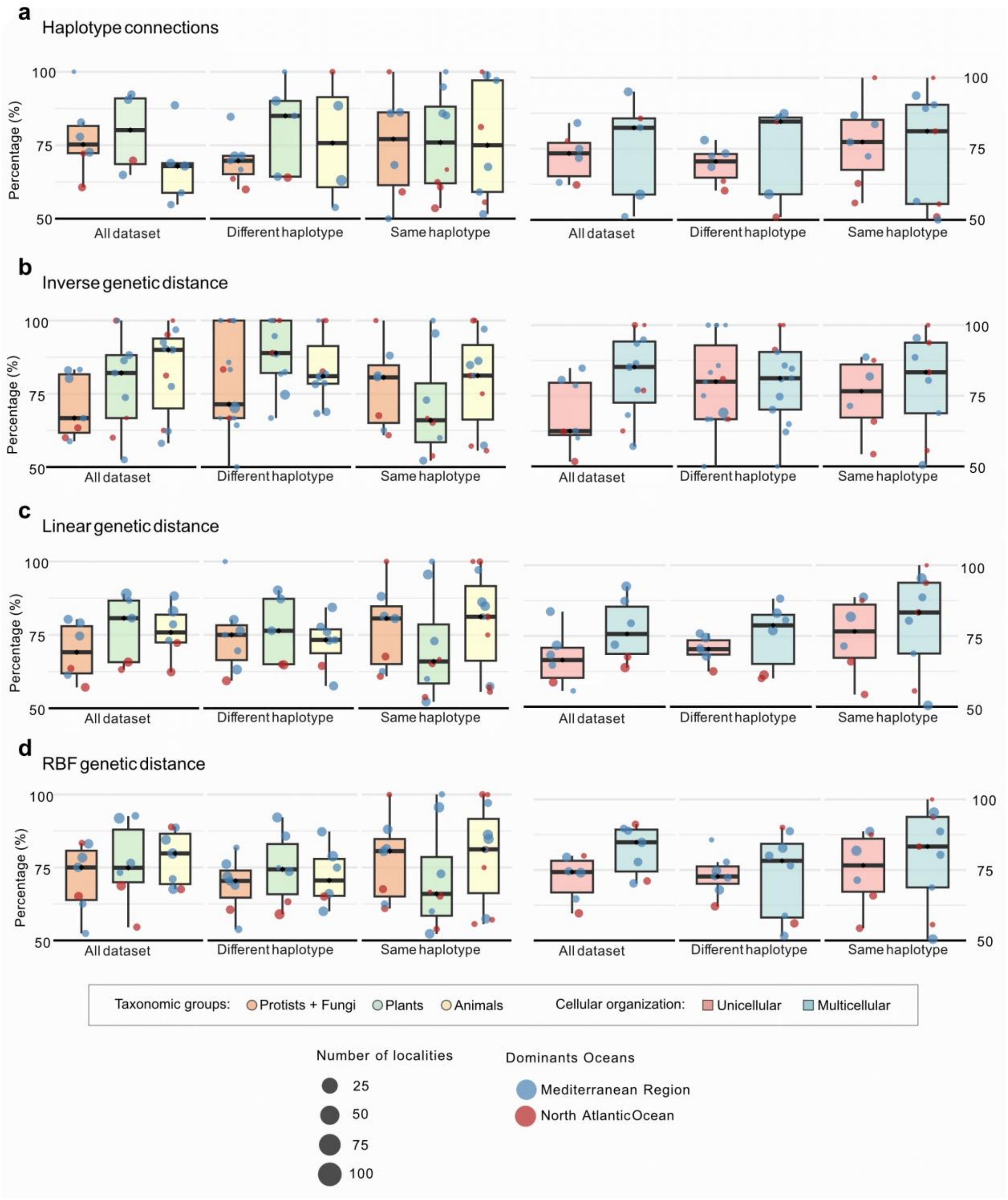


**Figure S12**. **Percentage of the dominant ocean within each locality cluster**. Boxplots showing the percentage of localities belonging to the dominant ocean within each cluster. Point colour represents the dominant ocean (Mediterranean Region in blue, North Atlantic Ocean in red), and point size indicates the number of localities composing each cluster. Within each panel, results for taxonomic groups (protists and fungi, plants and animals) are shown on the left, whereas results according to cellular organization (unicellular and multicellular taxa) are shown on the right. Results are presented for networks constructed using: **a**, haplotype-network connections only; **b**, inverse genetic-distance weighting; **c**, linear genetic-distance weighting; and **d**, an RBF (radial basis function) transformation of genetic distance.

**Table S1. Spearman's rank correlations among nucleotide diversity, relative abundance, and haplotype connectivity across broad taxonomic groups.** Correlations were calculated for the complete dataset (Global) and separately for plants, animals, and protists/fungi using metric values obtained for geographic clusters across informative OTUs. Relative abundance and haplotype connectivity were log-transformed before analysis. $\rho$ denotes Spearman's rank correlation coefficient, $P$ the associated two-tailed significance value, and $n$ the number of observations included in each analysis. Available from Zenodo as Table_S1.csv

**Table S2. Spearman's rank correlations among nucleotide diversity, relative abundance, and haplotype connectivity according to cellular organization.** Correlations were calculated for the complete dataset (Global) and separately for multicellular and unicellular taxa using metric values obtained for geographic clusters across informative OTUs. Relative abundance and haplotype connectivity were log-transformed before analysis. $\rho$ denotes Spearman's rank correlation coefficient, $P$ the associated two-tailed significance value, and $n$ the number of observations included in each analysis. Available from Zenodo as Table_S2.csv

**Table S3. Network modularity of locality clusters across taxonomic groups and cellular organization categories.** The table reports modularity values for locality networks generated using different weighting schemes. Groups include broad taxonomic categories (animals, plants and protists/fungi), and cellular organization categories (unicellular and multicellular taxa). Haplotype-connection types indicate whether networks were built using all haplotype connections ("all"), connections between the same haplotype ("same") or between different haplotypes ("diff"). Cluster indicates the number of detected locality clusters. Available from Zenodo as Table_S3.csv

**Table S4. Spatial autocorrelation of locality clusters and oceanic-region composition.** The table reports Moran's I values to evaluate spatial autocorrelation in locality-cluster structure using different weighting schemes. Groups include broad taxonomic categories (animals, plants and protists/fungi), and cellular organization categories (unicellular and multicellular taxa). Haplotype-connection types indicate whether networks were built using all haplotype connections ("all"), connections between same haplotype ("same") or between different haplotypes ("diff"). Moran's I values are associated with their significance levels (p), the number of detected locality clusters (Cluster), and the number of localities or observations assigned to each cluster (Freq). Mediterranean Region and North Atlantic Ocean indicate the percentage contribution of each oceanic region to the cluster. Available from Zenodo as Table_S4.csv

**Data S1. eKOI database ver. 2.0.** The database comprises 1,095,897 amplicon sequence variants (ASVs) and 362,781,685 reads obtained from 4,659 samples distributed across 1,097 unique localities. Each FASTA sequence header contains a unique eKOI ASV identifier followed by the sample identifiers of the localities in which the ASV was detected and its corresponding read abundance (reads) in each locality. Sample identifiers can be cross-referenced with the metadata provided in Data S2. Available from Zenodo as Data_S1.fasta

**Data S2. Metadata of eKOI 2.0 localities included in this study.** Locality metadata for the eKOI metabarcoding dataset 2.0 used in this study. The file includes sample identifiers (id_sample), source publications (paper), geographic coordinates, ecological realm (ecology), oceanic region and sea or marine subregion for 4,659 marine samples. These metadata were used to assign samples to geographic localities and oceanic regions for downstream biogeographic and spatial genetic analyses. Available from Zenodo as Data_S2.csv

**Data S3. Locality-cluster metrics used for abundance, diversity and connectivity distance-decay analyses.** This file contains locality-cluster-level metrics calculated for each OTU across eukaryotic phyla. For each OTU and locality cluster, the table reports taxonomic group, cellular organization, geographic coordinates, nucleotide diversity, maximum nucleotide diversity, total and log-transformed abundance, haplotype-network connectivity measured as the number of connections and log-connections, and geographic distances from the metric-specific centres of nucleotide diversity, abundance and connectivity. These data were used to fit distance-decay models and to compare spatial changes in abundance, intraspecific diversity and haplotype-network connectivity across phyla. Available from Zenodo as Data_S3.csv

**Data S4. Pairwise geographic and genetic distances classified by taxonomic group.** This file contains pairwise comparisons among locality clusters for each OTU, classified into broad taxonomic groups: animals, plants and protists/fungi. For each comparison, the table reports the source and target locality clusters, geographic coordinates, geographic distance in metres (m) and kilometres (km), genetic distance, haplotype-connection type, OTU identifier, haplotype information and phylum assignment. Geographic distances were calculated from locality-cluster coordinates and are provided using dots as decimal separators. Available from Zenodo as Data_S4.csv

**Data S5. Pairwise geographic and genetic distances classified by cellular organization.** This file contains pairwise comparisons among locality clusters for each OTU, classified according to cellular organization as unicellular or multicellular taxa. For each pairwise comparison, the table reports the source and target locality clusters, their geographic coordinates, geographic distance in metres (m) and kilometres (km), genetic distance, haplotype-connection type, OTU identifier, haplotype information and phylum assignment. Geographic distances were calculated from locality-cluster coordinates and are provided using dots as decimal separators. Available from Zenodo as Data_S5.csv
